# Cohesin mediates DNA loop extrusion and sister chromatid cohesion by distinct mechanisms

**DOI:** 10.1101/2022.09.23.509019

**Authors:** Kota Nagasaka, Iain F. Davidson, Roman R. Stocsits, Wen Tang, Gordana Wutz, Paul Betty, Melanie Panarotto, Gabriele Litos, Alexander Schleiffer, Daniel Wolfram Gerlich, Jan-Michael Peters

## Abstract

Cohesin connects CTCF binding sites and other genomic loci in *cis* to form chromatin loops, and replicated DNA molecules in *trans* to mediate sister chromatid cohesion. Whether cohesin uses distinct or related mechanisms to perform these functions is unknown. Here we describe a cohesin hinge mutant, which can extrude DNA into loops but is unable to mediate cohesion. Our results suggest that the latter defect arises during cohesion establishment. The observation that cohesin’s cohesion and loop extrusion activities can be separated indicates that cohesin uses distinct mechanisms to perform these two functions. Unexpectedly, the same hinge mutant can also not be stopped by CTCF boundaries as well as wildtype cohesin. This suggests that cohesion establishment and cohesin’s interaction with CTCF boundaries depend on related mechanisms and raises the possibility that both require transient hinge opening to entrap DNA inside the cohesin ring.

## Introduction

The structure, function and segregation of eukaryotic chromosomes depends on DNA-DNA interactions mediated by ‘structural maintenance of chromosomes’ (SMC) complexes (reviewed in Davidson and Peters, 2021; Yatskevich et al., 2019). The SMC complex cohesin forms two types of such interactions in eukaryotic interphase cells, *cis*-contacts between different loci on the same DNA molecule (Gassler et al., 2017; Hadjur et al., 2009; Nativio et al., 2009; Rao et al., 2014; Schwarzer et al., 2017; Wutz et al., 2017) and *trans*-contacts between replicated “sister” DNA molecules (Guacci et al., 1997; Michaelis et al., 1997). Cohesin mediated chromosomal *cis*-contacts exist in most cell types throughout interphase and lead to the formation of chromatin loops (Dixon et al., 2012; Nora et al., 2012; Rao et al., 2014), which have important roles in chromatin organization, gene regulation and recombination (Davidson and Peters, 2021). *Trans*-contacts between replicated DNA molecules are formed during S-phase in proliferating cells, persist until the onset of chromosome segregation in anaphase, and co-exist during this period with *cis*-contacts on the replicated DNA molecules. *Trans*-contacts result in sister chromatid cohesion, which enables DNA repair by homologous recombination between sister DNA molecules in G2-phase and the bi-orientation of chromosomes on the mitotic spindle in metaphase. This bi-orientation and the sister chromatid cohesion that it depends on are essential for chromosome segregation and cell proliferation.

The DNA-DNA contacts that form chromatin loops and cohesion could be of similar nature except for occuring in either *cis* or *trans*, in which case cohesin could use related mechanisms to create these contacts. However, several observations imply that cohesin interacts differently with DNA, depending on whether cohesin extrudes DNA or mediates cohesion. Experiments using mini-chromosomes in budding yeast indicate that cohesin generates cohesion by entrapping the replicated sister DNA molecules inside a ring structure that is formed by three of cohesin’s subunits. Two of these, called SMC1 and SMC3, form 50 nm-long anti-parallel coiled coils, which are connected to hinge domains at one and to ATP binding domains at their other ends (Haering et al., 2002). SMC1 and SMC3 form molecular rings by heterodimerizing via their hinge domains and by binding with their ATP binding domains to the N-terminal and C-terminal parts of a flexible “kleisin” subunit, called SCC1 (also known as RAD21 or Mcd1). Opening of the resulting SMC-kleisin rings (SK rings), possibly via transient dissociation of the hinge domains (Gruber et al., 2006; Collier and Nasmyth, 2022), is essential for entrapment of DNA and for cohesion (Haering et al., 2008).

In contrast, recent observations indicate that cohesin can form *cis*-contacts on DNA molecules without having to open its ring structure and might therefore perform this function without entrapping DNA inside the SK ring (Davidson et al., 2019). *In vitro*, cohesin forms these *cis*-contacts by binding to DNA and actively reeling it into loops, thereby bringing distant loci into proximity (Davidson et al., 2019; Golfier et al., 2020; Kim et al., 2019). This loop extrusion process can even occur on DNA molecules that are bound to particles that are bigger than cohesin’s ring size, supporting the hypothesis that cohesin forms chromosomal *cis*-contacts without having to entrap DNA inside its SK ring structure (Pradhan et al., 2021). These observations suggest that cohesin uses different mechanisms to mediate cohesion and to perform loop extrusion.

This possibility is also consistent with findings, which indicate that cohesin complexes that mediate cohesion (cohesive cohesin) and those that form *cis*-contacts (loop extruding cohesin) differ in their post-translational modifications, interactions with other proteins, and in their positions on genomic DNA (reviewed in Davidson and Peters, 2021). Cohesin needs to be acetylated on two lysine residues on SMC3’s ATP binding domain (Rolef Ben-Shahar et al., 2008; Rowland et al., 2009; Unal et al., 2008) and requires PDS5 proteins (Hartman et al., 2000; Panizza et al., 2000) and sororin (Rankin et al., 2005; Schmitz et al., 2007; Nishiyama et al., 2010) to maintain cohesion but can extrude DNA without these modifications and proteins (Davidson et al., 2019; Kim et al., 2019). Instead, cohesin can only perform loop extrusion in the presence of a PDS5-related protein called NIPBL (called Scc2 in budding yeast) (Davidson et al., 2019; Kim et al., 2019). Both PDS5 proteins and NIPBL are composed of huntingtin-elongation factor 3-protein phosphatase 2A-Tor1 (HEAT) repeats and associate in a mutually exclusive manner with SCC1 (Kikuchi et al., 2016; Petela et al., 2018). Of these ‘HEAT repeat subunits associated with kleisins’ (HAWKs; Wells et al., 2017) only NIPBL can support loop extrusion, whereas PDS5 proteins cannot (Davidson et al., 2019). Loop extruding and cohesive cohesin might therefore differ with respect to which of these HAWK subunits they interact with. A third HAWK subunit is more stably associated with cohesin than NIPBL and PDS5 proteins and is required for both cohesion (Roig et al., 2014) and loop extrusion (Davidson et al., 2019). Mammalian somatic cells contain two paralogs of this subunit, called STAG1 and STAG2 (Losada et al., 2000; Sumara et al., 2000), which are orthologs of Scc3 in budding yeast (Toth et al., 1999).

In cells, loop extruding cohesin complexes are constrained in their movements at specific genomic sites (Fudenberg et al., 2016; Sanborn et al., 2015; Banigan et al., 2021; Dequeker et al., 2022). The best characterized of these boundary elements are DNA sequences associated with the zinc finger protein CTCF (Parelho et al., 2008; Wendt et al., 2008) which is required for the formation of boundaries between ‘topologically associating domains’ (Nora et al., 2012; Wutz et al., 2017). These TADs emerge in analyses of large cell populations but represent collections of chromatin loops of which only subsets are present in individual cells (Bintu et al., 2018; Finn et al., 2019; Flyamer et al., 2017; Gabriele et al., 2022; Beckwith et al., 2021). Loops inside TADs might represent nascent loops that are in the process of being extruded, whereas loops that connect TAD boundaries are thought to be actively ‘anchored’ there by CTCF. CTCF can bind cohesin and stabilize it on chromatin by protecting it from WAPL (Li et al., 2020; Wutz et al., 2020), a protein which can release cohesin from DNA and thereby limit its residence time on chromatin (Kueng et al., 2006). However, this effect may not be CTCF’s only role in establishing loop extrusion boundaries, since cohesin still accumulates at genomic sites when these are bound by a CTCF mutant, which cannot bind cohesin (Li et al., 2020).

Here we report the identification of a cohesin hinge mutant, which separates cohesin’s DNA loop extrusion activity from its function in sister chromatid cohesion. Characterization of this separation-of-function allele in human cells (SMC1^3D^) and single-molecule DNA loop extrusion assays indicate that this cohesin mutant can extrude DNA into loops but cannot respond as well as wild-type cohesin to CTCF boundaries and is unable to mediate cohesion. Our results suggest that the latter defect arises early during cohesion establishment and is not simply caused by precocious release of this mutant from chromatin after cohesion has been established. Our finding that both cohesion establishment and cohesin’s interaction with CTCF boundaries require cohesin’s hinge suggests that these two processes depend on related mechanisms. In line with previous proposals (Gruber et al., 2006; Li and Dekker, 2021; Collier and Nasmyth, 2022), we speculate that both functions could depend on entrapment of DNA inside cohesin’s SK ring via transient opening of the hinge. By contrast, our observation that cohesin’s cohesion and loop extrusion activities can be genetically separated indicates that cohesin uses at least partially distinct mechanisms to perform these functions.

## Results

### A hinge mutant of human cohesin is defective in sister chromatid cohesion, stable association with chromatin in G2-phase and sororin binding

To test whether cohesin uses distinct or related mechanisms for loop extrusion and sister chromatid cohesion we searched for cohesin mutants that are deficient in one but proficient in the other function. Versions of budding yeast cohesin in which basic amino acid residues in the inner pore of the hinge were replaced by acidic and neutral residues have been proposed to represent such separation-of-function mutants, since at least one of these inner-pore hinge mutants cannot support cohesion but still enables the formation of rDNA loops, which can be detected microscopically (Srinivasan et al., 2018). However, it is unknown whether rDNA loops are formed by loop extrusion and loop extrusion mediated by budding yeast cohesin has not been reconstituted in vitro yet (Ryu et al., 2021). We therefore searched for equivalent mutations in human cohesin, for which assays have been developed in which loop extrusion can be observed (Davidson et al., 2019; Kim et al., 2019), to test whether cohesin’s cohesion and extrusion functions can be separated.

For this purpose, we generated a human cell line, in which endogenous SMC1 can be replaced with ectopically expressed versions of SMC1 and in which SCC1 can be simultaneously visualized with a fluorescent dye. We modified all alleles of endogenous SMC1 and SCC1 in Hela cells by genome engineering to fuse AID-mKate2 and Halo-P2A-Tir1 to the C-termini of SMC1 and SCC1, respectively (Figures S1A-C). The SCC1-Halo and SMC1-AID-mKate2 proteins encoded by these alleles were functional in supporting chromosomal association of cohesin (Figure 1A, lanes 1 and 4) and cell proliferation (Figure 1B, left). Addition of auxin led to degradation of SMC1-AID-mKate2, disappearance of other cohesin subunits from chromatin fractions (Figure 1A, lanes 2 and 5) and loss of cell proliferation (Figures 1B, ‘+aux’), indicating efficient depletion of cohesin.

**Figure 1.**
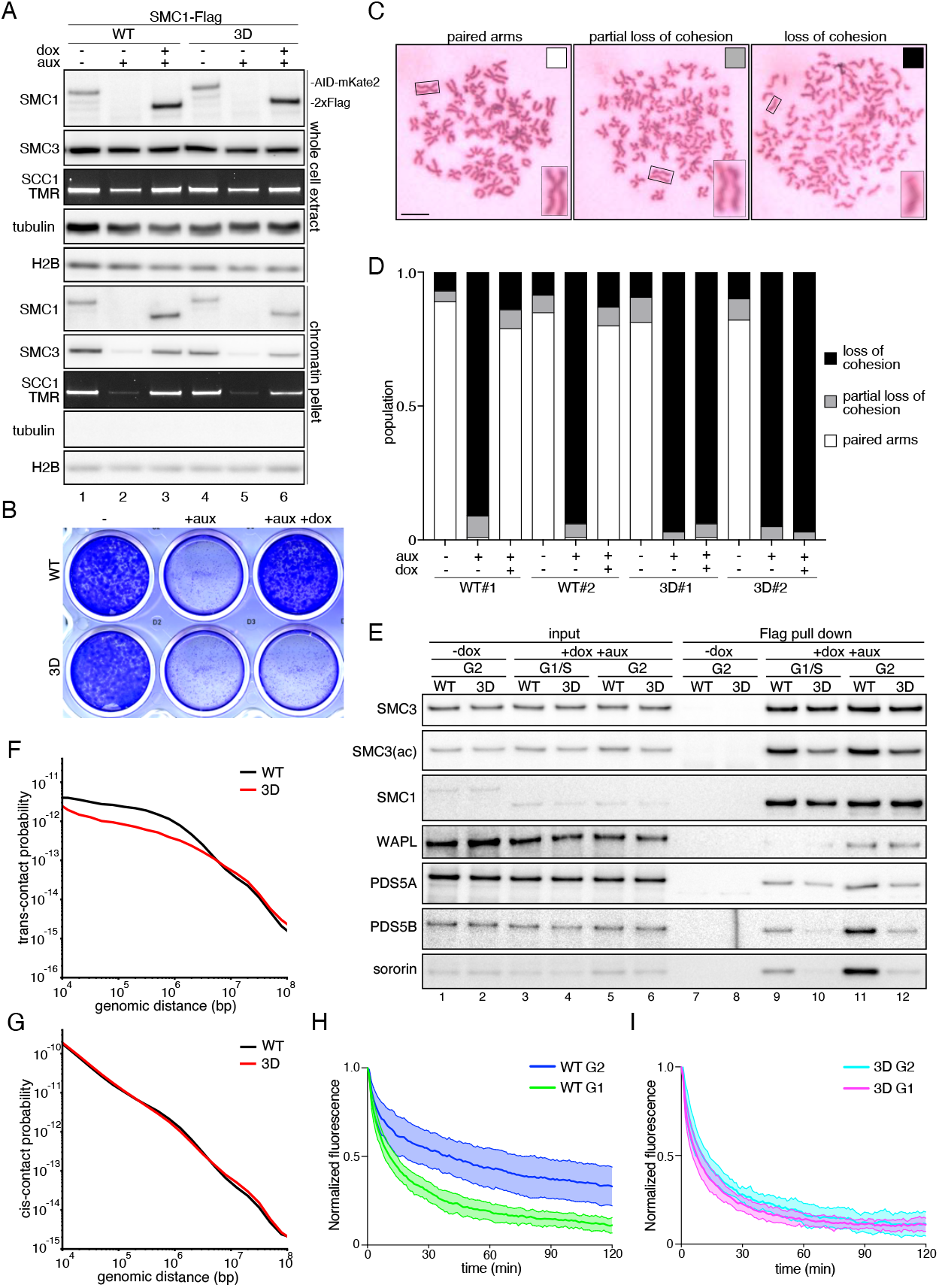
Cohesin^3D^ is defective in sister chromatid cohesion, stable chromatin binding in G2-phase and sororin recruitment. (A) Replacement of endogenous SMC1 proteins with Flag-tagged SMC1^WT^ and SMC1^3D^ proteins in HeLa cells. Whole cell extracts and chromatin fractions of the resulting cohesin^WT^ and cohesin^3D^ cells were analyzed by immunoblotting as indicated. (B) Images of a plate containing cohesin^WT^ and cohesin^3D^ cells, visualized by Crystal Violet staining for the presence of viable cells after addition of auxin for 5 days and subsequent addition of doxycycline for 48 hours. (C) Representative images of mitotic chromosomes from cells expressing cohesin^WT^ and cohesin^3D^, analyzed by spreading and Giemsa staining. Scale bar,10 μm. (D) Quantification of cohesion phenotypes shown in (C). (E) Immunoblot analysis of SMC1^WT^ and SMC1^3D^ immunoprecipitates obtained with Flag antibodies. (F, G) Average contact probability over different genomic distances for *trans* (H) and *cis* (I) contacts obtained by scs-Hi-C from cohesin^WT^ and cohesin^3D^ cells in G2 phase. (H, I) Normalized recovery kinetics of SCC1-Halo^TMR^ signals after photobleaching of cohesin^WT^ (H) and cohesin^3D^ (I) cells in G1 and G2-phase (for representative images see Figure S1H). Scale bar, 10 μm.

We ectopically expressed SMC1 mutants under control of a doxycycline inducible promoter in these cells. Either two or three lysine residues in SMC1’s hinge were replaced by aspartic acid (SMC 1^K540D/K637D^, SMC 1^K540D/K648D^, SMC1^K637D/K648D^ and SMC1^K540D/K637D/K648D^; we refer to the triple mutant as SMC1^3D^; Figure S1D-F). Ectopically expressed wild-type SMC1-AID-mKate2 (SMC1^WT^) and the double mutants, but not SMC1^3D^ were able to support cell proliferation when their expression was induced before endogenous SMC1 degradation had been induced (Figures 1B, ‘+aux +dox’ and S1G). This indicates that SMC1^3D^ abrogates an essential cohesin function in human cells, as do the related hinge mutants in yeast (Srinivasan et al., 2018).

To address whether SMC1^3D^ can support cohesion, we analyzed the morphology of mitotic chromosomes in chromosome spreads (Figure 1C and D). Addition of auxin for 3 hours caused extensive separation of sister chromatids, indicating loss of cohesion. This defect could largely be prevented by inducing expression of SMC1^WT^ but not by SMC1^3D^ 48 hours before addition of auxin (Figure 1C and D). SMC1^3D^ associated with chromatin and enabled other cohesin subunits to do so, although to a lesser extent than SMC1^WT^ (Figure 1A, lanes 3 and 6 ‘chromatin pellet’). In immunoprecipitation experiments SMC1^3D^ co-precipitated other cohesin core subunits similarly well as SMC1^WT^ (Figure 1E, lanes 9-12). These results indicate that SMC1^3D^ assembles into cohesin complexes that can bind to DNA but are defective in cohesion.

To analyze whether the cohesion defect caused by SMC1^3D^ occurs only in mitosis or already during G2-phase, we performed sister chromatid sensitive Hi-C (scsHi-C) experiments, with which DNA-DNA *cis* and *trans* contacts can be detected (Mitter et al., 2020). These experiments revealed a global reduction of *trans*-contacts in cells expressing SMC1^3D^ compared to cells expressing SMC1^WT^ (Figure 1F), whereas the number of *cis* contacts and the genomic distances bridged by these were similar in the presence of SMC1^WT^ and SMC1^3D^ (Figure 1G). These results indicate that the cohesion defects caused by SMC1^3D^ exist already during G2-phase.

This finding predicts that in G2-phase cohesin containing SMC1^3D^ (cohesin^3D^) should bind to chromatin less stably than wildtype cohesin, most of which associates with chromatin only for minutes during G1 but about half of which binds chromatin for hours during G2 (Gerlich et al., 2006). For this purpose, we labeled endogenous SCC1-Halo with TMR and monitored the recovery kinetics of SCC1-halo^TMR^ after photobleaching. We performed these measurements in the presence of excess unlabeled Halo tag ligand to avoid effects of newly synthesized SCC1-Halo (Rhodes et al., 2017). These pulse-chase inverse FRAP (pciFRAP) experiments revealed that in G1-phase, cohesin containing SMC1^WT^ (cohesin^WT^) bound chromatin with a residence time of 8-20 min, whereas in G2-phase 60% of these complexes bound with a residence time of 4-7 hours (Figure 1H and S1H-K), consistent with previous reports (Gerlich et al., 2006). In contrast, pciFRAP curves of cohesin^3D^ showed little, if any increase in stably bound fractions in G2-phase (Figure 1I). By contrast, the pciFRAP curves from cohesin^3D^ in G1-phase showed only slightly faster recovery kinetics than those of cohesin^WT^ (Figure 1I and S1L), indicating that SMC1^3D^ had only little effect on the chromosomal association of cohesin prior to DNA replication. Cohesin^3D^ is therefore unable to stably associate with chromatin in G2-phase.

The long residence time of cohesive cohesin in G2-phase and the ability of these complexes to maintain cohesion from S-phase until mitosis depends on sororin (Ladurner et al., 2016; Nishiyama et al., 2010; Rankin et al., 2005; Schmitz et al., 2007). Sororin interacts with cohesive cohesin in a manner that depends on SMC3 acetylation (Lafont et al., 2010; Nishiyama et al., 2010), which in turn depends on PDS5 proteins (Chan et al., 2013; Minamino et al., 2015; Vaur et al., 2012). We therefore analyzed whether cohesin^3D^ becomes acetylated and can associate with PDS5 proteins and sororin. Immunoblot analyses of immunoprecipitated cohesin complexes indicated that cohesin^3D^ associated with much less PDS5A, PDS5B and even lesser sororin in G2 phase than cohesin^WT^. In contrast, SMC3 acetylation levels were only moderately reduced in cohesin^3D^ (Figure 1E, lanes 9-12). Taken together, these results show that cohesin^3D^ is unable to associate properly with sororin and PDS5 proteins, cannot mediate cohesion, and is unable to associate stably with chromatin in G2-phase.

### WAPL depletion stabilizes cohesin^3D^ on chromatin but does not prevent the cohesion and proliferation defects caused by SMC1^3D^

Sororin is essential for cohesion in the presence of WAPL but not in its absence (Nishiyama et al., 2010). This and other observations (Ladurner et al., 2016) indicate that sororin maintains cohesion by preventing precocious release of cohesive cohesin from chromatin by WAPL but is not required for cohesion *per se*. If the inability of cohesin^3D^ to mediate cohesion was merely due to its failure to bind sororin, WAPL depletion should therefore prevent cohesion loss in cells expressing cohesin^3D^.

-To test this possibility, we modified all *WAPL* alleles in the cell lines, in which endogenous SMC1 can be replaced with SMC1^WT^ or SMC1^3D^, to enable inducible WAPL degradation. For this purpose, we engineered the *WAPL* alleles to encode fusions with N-terminal FKBP12^F36V^ tags (Figure S1A, S2A and S2B), which mediate protein degradation upon addition of dTAG (Nabet et al., 2018). We induced expression of SMC1^WT^ or SMC1^3D^ in these cells, arrested them at the G1-S-phase transition by double-thymidine arrest, released them in the presence or absence of auxin and dTAG to induce degradation of endogenous SMC1 and WAPL, respectively (Figure 2A), and analyzed sister chromatid cohesion in the subsequent mitosis by chromosome spreading (Figure 2B and C and S2C). As a control, we depleted sororin by RNAi in some of these cells. In cells expressing ectopically expressed SMC1^WT^, WAPL degradation resulted in chromosomes with tightly aligned ‘over-cohesed’ arms, consistent with previous reports (Gandhi et al., 2006; Kueng et al., 2006). In contrast, WAPL degradation did not prevent loss of cohesion in cohesin^3D^ cells. This lack of rescue was not due to insufficient WAPL degradation since cohesion loss induced by sororin depletion could in most cells be prevented by WAPL degradation (Figure 2B and C). In agreement with the above observations, WAPL degradation restored cell proliferation in sororin-depleted cells (Figure S2D) but not in cohesin^3D^ cells (Figure 2D). The cohesion defects caused by cohesin^3D^ are therefore not merely due to reduced interactions of this mutant with sororin.

**Figure 2.**
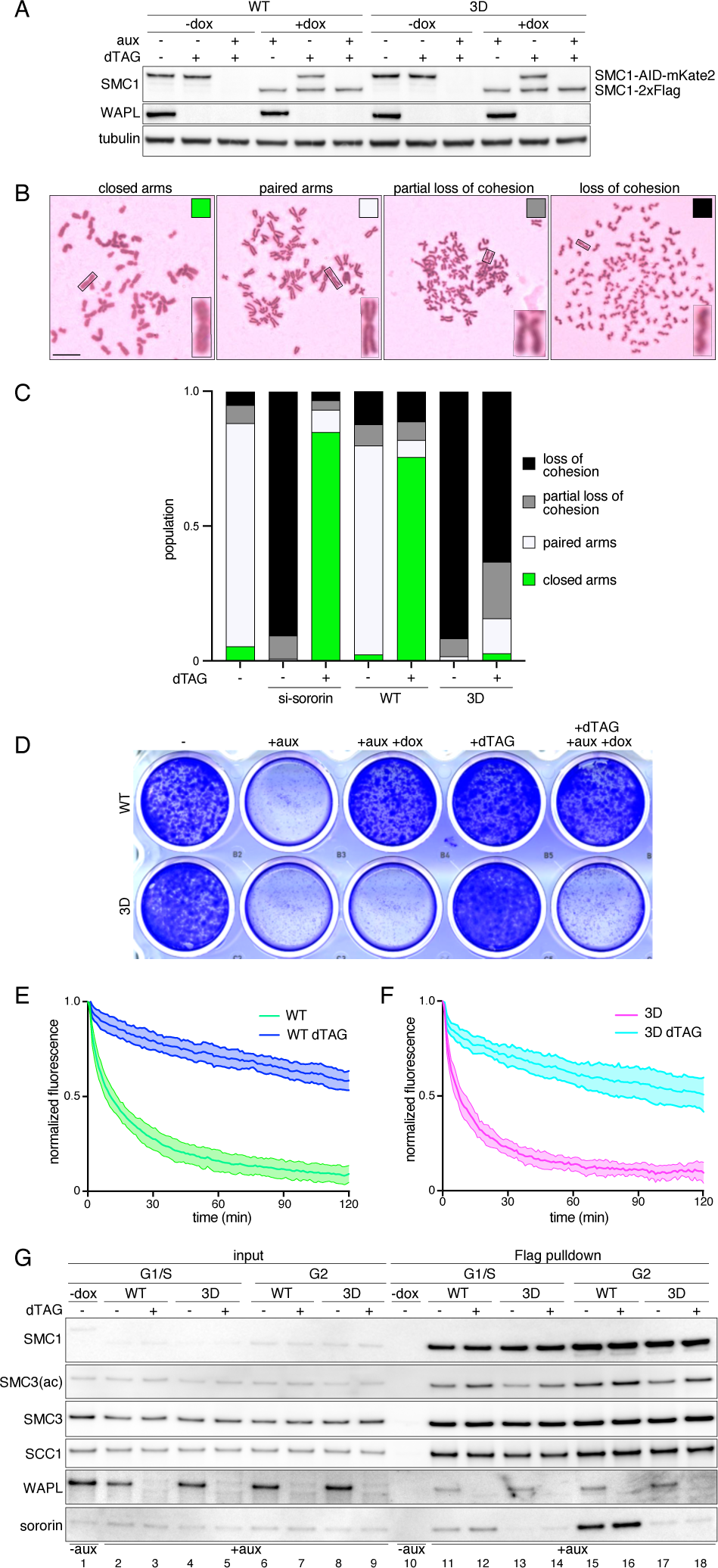
WAPL degradation stabilizes cohesin^3D^ on chromatin but fails to restore cohesion. (A) Degradation of WAPL via the dTAG system in cohesin^WT^ and cohesin^3D^ cells. Whole cell extracts were analyzed by immunoblotting as indicated. (B) Representative images of mitotic chromosomes from cells expressing cohesin^3D^, analyzed by spreading and Giemsa staining (C) Quantification of the cohesion phenotypes shown in (B) (for additional analyses see Figure S2C). (D) Images of plates containing cohesin^WT^ and cohesin^3D^ cells, analyzed by Crystal Violet staining for the presence of viable cells after addition of auxin for 5 days and subsequent addition of dTAG7 for 5 hours and doxycycline for 48 hours. (E, F) Normalized recovery kinetics of SCC1-Halo^TMR^ signals after photobleaching of cohesin^WT^ (E) and cohesin^3D^ (F) cells in G1-phase following degradation of WAPL. (G) Immunoblot analysis of SMC1^WT^ and SMC1^3D^ immunoprecipitates obtained with Flag antibodies from cells in which WAPL had been degraded or not, as indicated.

Instead, cohesin complexes containing SMC1^3D^ could be unable to maintain cohesion because they cannot stably associate with chromatin at all and would therefore be precociously released from chromatin in a WAPL independent manner. If so, cohesin^3D^ would not stably associate with chromatin even in the absence of WAPL. To test this possibility, we induced WAPL degradation by dTAG and measured cohesin’s chromatin residence time via fluorescently labeled SCC1-Halo^TMR^ in cohesin^WT^ and cohesin^3D^ cells by pciFRAP. However, WAPL degradation similary increased the residence time of cohesin^WT^ and cohesin^3D^ to 4-9 hours with 90% of cohesin being stably bound to chromatin in G1-phase (Figure 2E and F and S2E-I). Also in G2-phase, where cohesin^3D^ does not stably bind to chromatin (Figure 1I), WAPL depletion increased the chromatin residence time almost to the one observed for cohesin^WT^ (Figure S2G and H). Similar observations were made in mitotic cells, in which most cohesin is released from chromosome arms in prophase and prometaphase (Waizenegger et al., 2000) unless WAPL is depleted (Gandhi et al., 2006; Kueng et al., 2006). Live cell imaging of mitotic cells expressing SCC1-Halo^TMR^ revealed that not only cohesin^WT^ but also cohesin^3D^ remained associated with chromosome arms if WAPL had been degraded, whereas in WAPL’s presence cohesin^WT^ and cohesin^3D^ were similarly dispersed throughout the cytoplasm after nuclear envelope breakdown, with only weak signals of cohesin^WT^, but not cohesin^3D^, remaining on centromeres and chromosome (Figure S2J).

Cohesin^3D^ can therefore stably associate with chromatin in the absence of WAPL. Even under these conditions, sororin did not co-immunoprecipitate with cohesin^3D^, even though WAPL depletion increased SMC3 acetylation on cohesin^3D^ (Figure 2G, compare lanes 16 and 18). These observations indicate that the inability of cohesin^3D^ to mediate cohesion is not caused by a principal disability in stable chromatin binding. As expected, these results also show that stabilization of cohesin on chromatin and SMC3 acetylation are not sufficient to generate cohesion and sororin-cohesin interactions.

### Cohesin^3D^ is able to extrude DNA into loops *in vitro*

To test whether the SMC1^3D^ mutant also affects loop extrusion, we generated and purified recombinant versions of cohesin^WT^ and cohesin^3D^ (Figure 3A) and analyzed their ability to extrude DNA into loops in real time by TIRF microscopy. We also measured the ATPase activities of these complexes, since cohesin’s ability to bind and hydrolyze ATP is essential for its loop extrusion activity (Davidson et al., 2019; Kim et al., 2019). We found that the ATPase activity of cohesin^3D^ was slightly increased (Figure 3B), whereas the frequency and to a lesser extent also the rates with which cohesin^3D^ extruded DNA molecules were reduced compared to those observed for cohesin^WT^ (Figure 3C-E). However, cohesin^3D^ complexes could clearly perform loop extrusion, even though they are unable to mediate cohesion.

**Figure 3.**
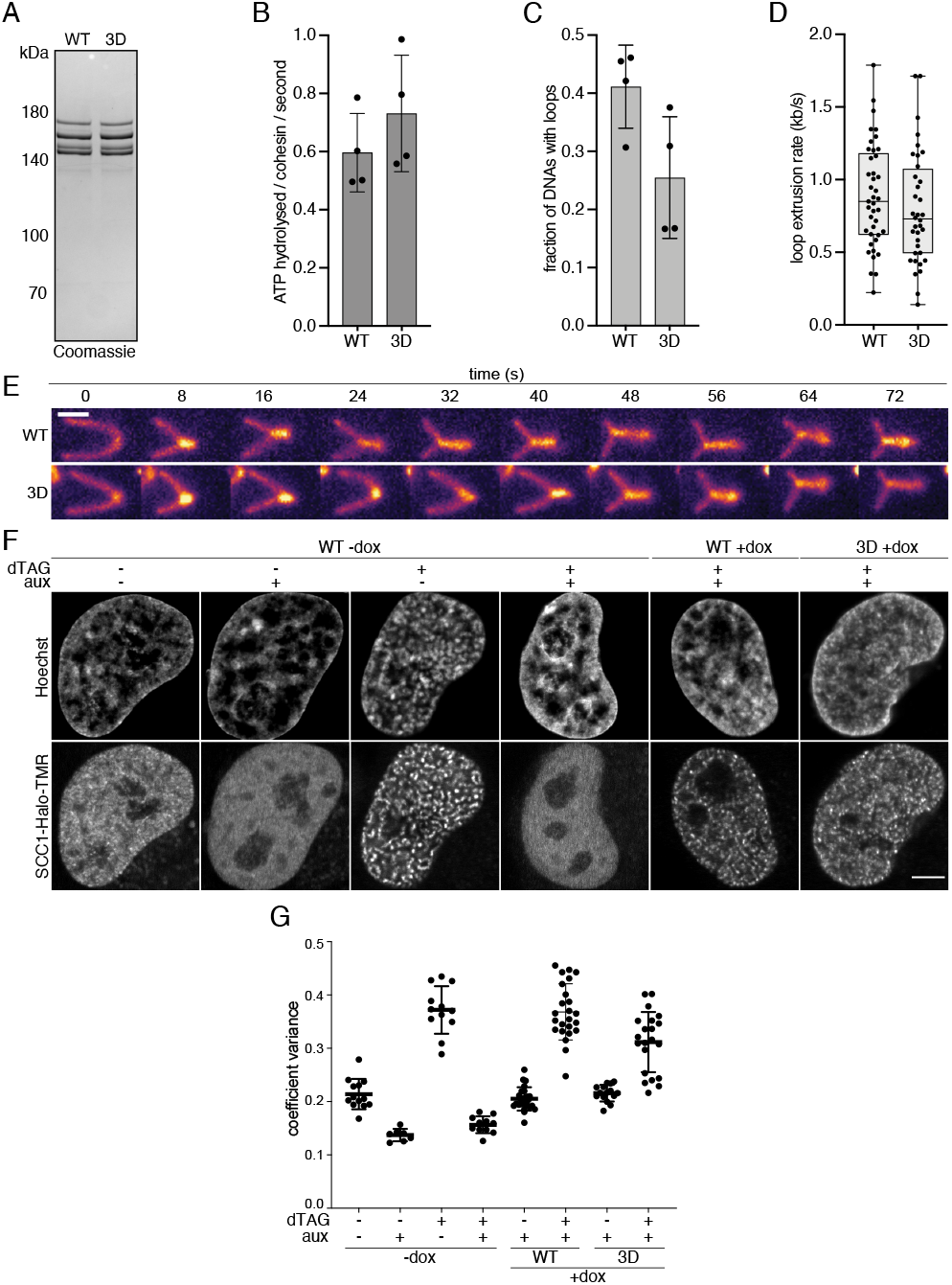
Cohesin^3D^ can perform DNA loop extrusion *in vitro* and form vermicelli in WAPL depleted cells. (A) Coomassie staining of recombinant cohesin^WT^ and cohesin^3D^ purified from insect cells and separated by SDS-PAGE. (B) ATP hydrolysis rates of cohesin^WT^ and cohesin^3D^ in the presence of DNA and NIBPL-MAU2. The error bars correspond to mean ± SD of four independent experiments. (C) Frequencies of DNA extrusion events catalyzed by cohesin^WT^ and cohesin^3D^ in the presence of NIPBL and ATP. (D) Rates of loop extrusion from individual events. Median and quartiles are shown. (E) Representative images of loop extrusion events catalyzed by cohesin^WT^ and cohesin^3D^. Scale bar,2 μm. (F) Representative live cell images of SCC1-Halo^TMR^ in cells from which WAPL had been depleted or not, as indicated. DNA was stained with Hoechst. Scale bar, 5 μm. (F) Quantification of vermicelli by measuring the coefficient variance of SCC1-Halo^TMR^ fluorescence intensity

### Cohesin^3D^ is able to form vermicelli and TADs in cells

To test whether cohesin^3D^ can also form chromatin loops in cells, we first analyzed the intra-nuclear distribution of cohesin in WAPL depleted cells by fluorescence microscopy. WAPL depletion causes chromatin compaction and the accumulation of cohesin in axial chromosomal domains, called vermicelli (Tedeschi et al., 2013). It has been suggested that the accumulation of cohesin in these regions reflects an extended loop extrusion activity of cohesin (Fudenberg et al., 2016; Haarhuis et al., 2017; Wutz et al., 2017). We therefore analyzed whether cohesin^3D^ can accumulate in vermicelli. Live cell imaging of SCC1-Halo^TMR^ showed that WAPL degradation induces vermicelli formation in cohesin^3D^ cells, although to a lesser extent than in cohesin^WT^ cells (Figure 3F). Quantification of these results by measuring the variance of SCC1-Halo^TMR^ signals within nuclei confirmed these observations (Figure 3G). These results suggest that cohesin^3D^ can generate chromatin loops also in cells.

To test this possibility more directly, we synchronized cells at the G1-S-phase transition and used Hi-C to analyze chromosomal *cis*-interactions. When cell populations are analyzed by this technique, TADs appear in Hi-C matrices as pyramid-shaped structures, which are composed of numerous *cis*-interactions. Different sub-sets of these exist in individual cells, presumably representing nascent chromatin loops that are in the process of being extruded. TAD patterns emerge from these asynchronous loops only when cis-interactions from many cells are super-imposed (Flyamer et al., 2017; Gabriele et al., 2022; Beckwith et al., 2021). By contrast, cis-interactions that occur simultaneously in many cells appear in Hi-C matrices as dots which are either located at the apices of TADs and are therefore called ‘corner peaks’ or are found at TAD boundaries (Rao et al., 2014). We collectively refer to these dots as ‘Hi-C peaks’.

In cells ectopically expressing SMC1^WT^ or SMC1^3D^, TADs could be detected in similar numbers (Figure 4A and B), sizes (Figure 4C) and with comparable insulation scores (Figure 4D), although in cohesin^3D^ cells the TAD signals were weaker (Figure 4A) and contacts in the 300 Kb-3 Mb range less frequent (Figure 4E) than in cohesin^WT^ cells. A and B compartments, long range interactions that correspond to euchromatic and heterochromatic regions, could be detected similarly well in cohesin^WT^ and cohesin^3D^ cells (Figure S3A).

**Figure 4.**
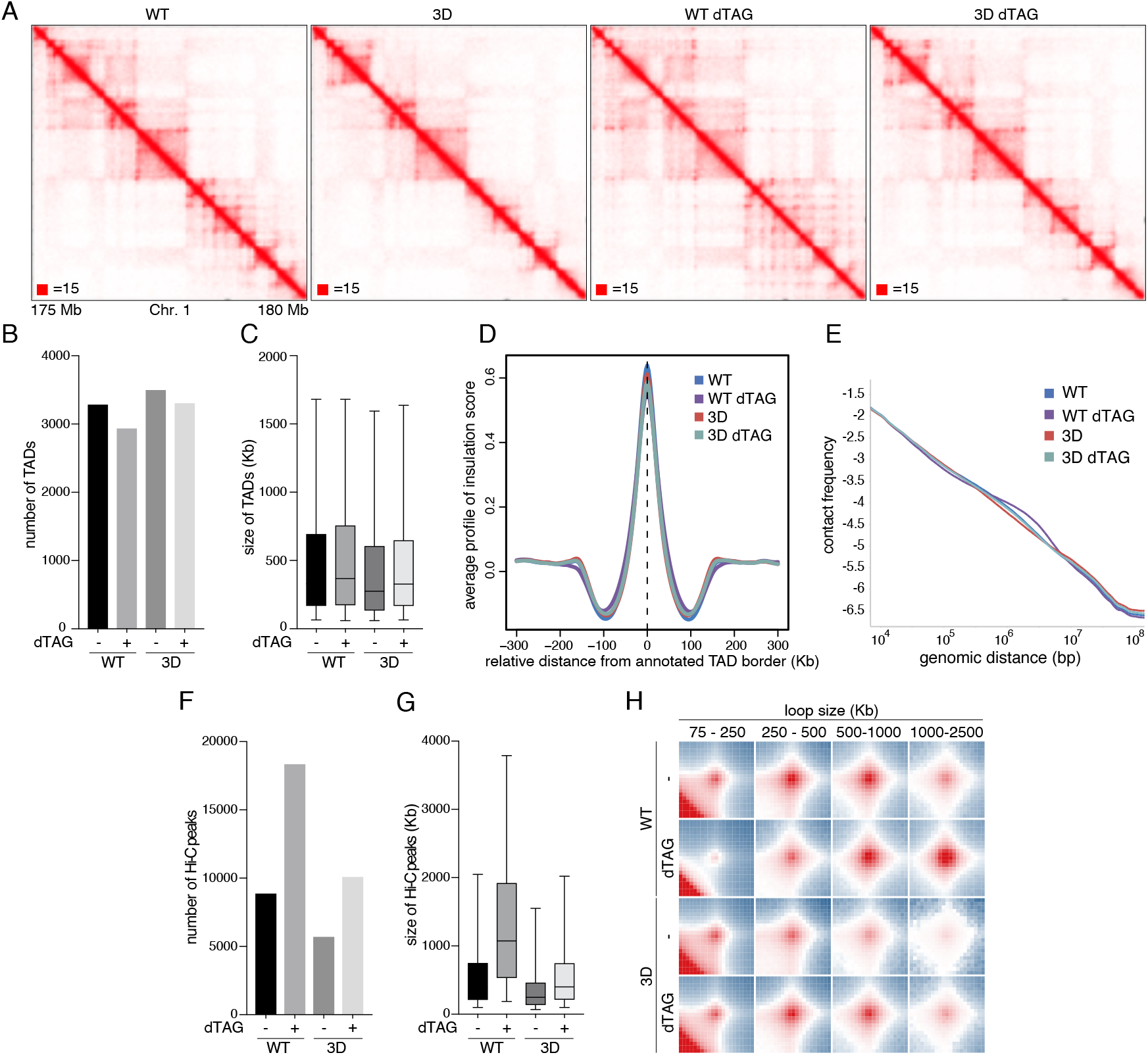
Cohesin^3D^ is able to form TADs, but not CTCF-anchored Hi-C peaks. (A) Balanced (Knight-Ruiz) normalized Hi-C contact matrices of chromosome 1 (175-180 Mb) of cohesin^WT^ and cohesin^3D^ cells from which WAPL had been depleted or not, as indicated. (B, C) Quantification of numbers (B) and sizes (C) of TADs. (D) Average insulation score around TAD boundaries in cohesin^WT^ and cohesin^3D^ cells. (E) Overall scaling of Hi-C contact frequency. (F, G) Quantification of number (F) and size (G) of Hi-C peaks. (H) Aggregated Hi-C peak analysis for chromatin loops of different lengths.

By contrast, the number of Hi-C peaks was greatly reduced in cohesin^3D^ cells (5,694 peaks) compared to cohesin^WT^ cells (8,862 peaks; Figure 4F). Aggregate peak analyses showed that on average the Hi-C peaks represented shorter *cis*-interactions in cohesin^3D^ than in cohesin^WT^ cells (Figure 4G and H). To test whether these differences could be caused by an increased sensitivity of cohesin^3D^ to WAPL, we analyzed chromosomal *cis*-interactions in cohesin^WT^ and cohesin^3D^ cells in which WAPL had been degraded (Figure 4A, right panels). WAPL depletion increased the number of Hi-C peaks in cohesin^WT^ (18,335 peaks) and to a lesser extent also in cohesin^3D^ cells (10,084 peaks; Figure 4F). As a result, the number of Hi-C peaks that could be detected in WAPL depleted cohesin^3D^ cells was similar to the peak number in cohesin^WT^ cells from which WAPL had not been depleted. These results suggest that increased sensitivity of cohesin^3D^ to WAPL can partially but not fully explain why this cohesin mutant forms fewer Hi-C peaks. Likewise, the median length of *cis*-interactions was increased in both cases but also in this case the length of *cis*-interactions observed in WAPL depleted cohesin^3D^ cells only reached a value observed in cohesin^WT^ cells in the presence of WAPL (Figure 4G and H). These results show that cohesin^3D^ can generate chromatin loops in cells, although not to the same extent as cohesin^WT^. Cohesin’s ability to extrude DNA does therefore not depend on its ability to mediate cohesion.

### In WAPL depleted cells, cohesin^3D^ accumulates in cohesin islands

To analyze whether SMC1^3D^ affects the genomic distribution of cohesin, we performed calibrated chromatin immunoprecipitation sequencing (ChIP-Seq) using Flag antibodies to isolate SMC1^WT^ and SMC1^3D^. For normalization, we spiked mouse embryonic fibroblasts (MEFs) expressing an unrelated Flag-tagged chromatin protein into all samples (Figure S4A).

We used -dox cells as an input track for peak calling and identified 29,439 SMC1-Flag specific ChIP-Seq peaks in cohesin^WT^ cells (Figures 5A and B and S4C). As expected, most SMC1^WT^ peaks (90%) overlapped with CTCF (Figure 5B-E). The numbers and positions of SMC1^WT^ peaks are comparable to those of peaks previously identified with SMC1 antibodies in HeLa cells (38,857 sites, 91.1 % overlap with cohesin^WT^; Figure S4B), indicating that cohesin containing ectopically expressed SMC1^WT^ behaves largely like endogenous cohesin with respect to its genomic localization.

**Figure 5.**
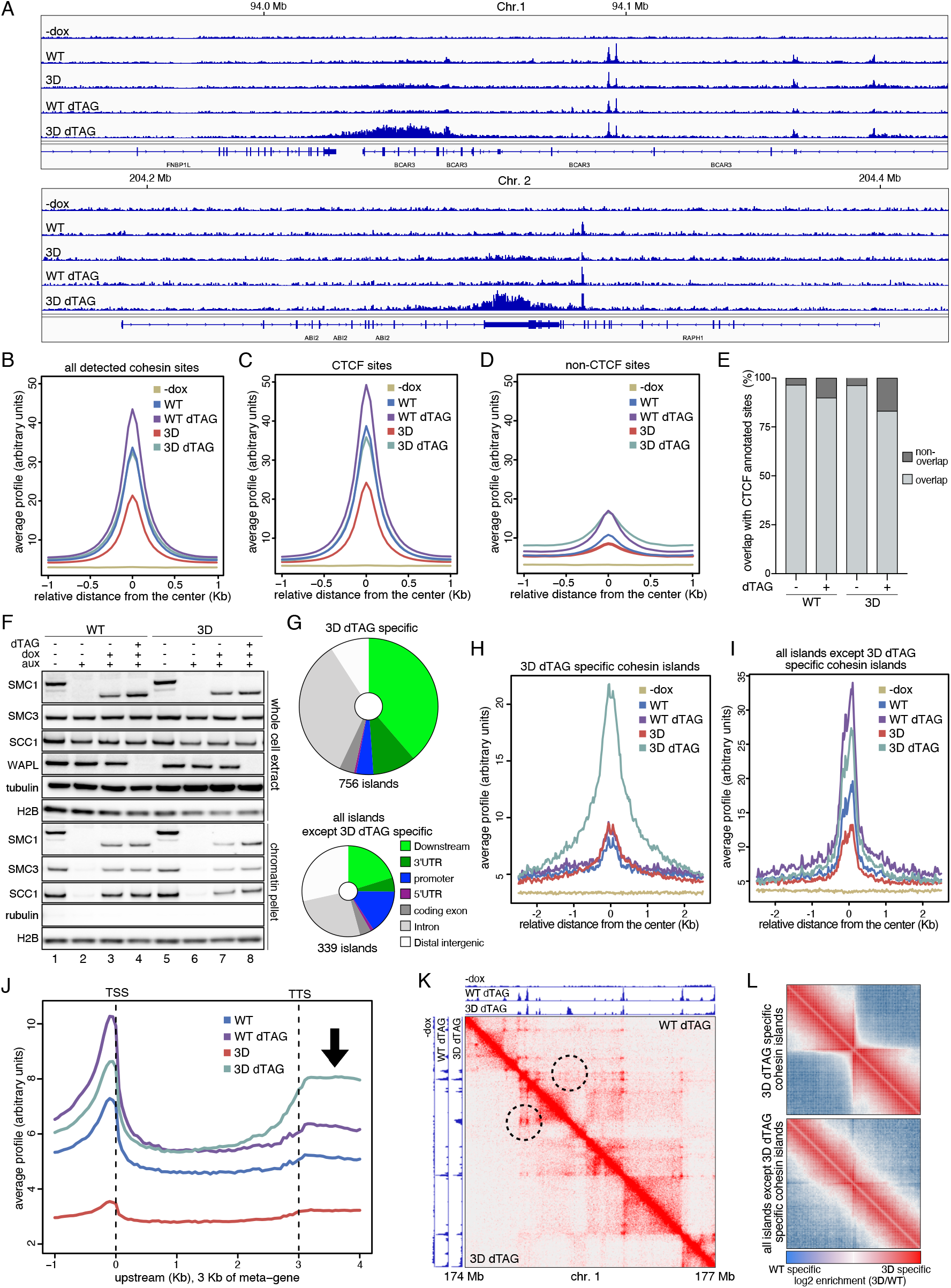
WAPL depletion is sufficient for accumulation of cohesin^3D^ in cohesin islands. (A) Calibrated ChIP-Seq profiles of SMC1^WT^ and SMC1^3D^ obtained with Flag antibodies from cells in which WAPL had been depleted or not, as indicated. (B-D) Average ChIP-Seq site profile at all cohesin sites (B), CTCF sites (C) and non-CTCF sites (D). (E) Fraction of SMC1^WT^ and SMC1^3D^ peaks at CTCF sites or at non-CTCF sites, obtained under the indicated conditions. (F) Immunoblot analysis of whole cell extracts and chromatin pellets from cohesin^WT^ and cohesin^3D^ cells from which WAPL had been depleted, as indicated. (G) Genomic localization of cohesin islands observed after WAPL depletion only in cohesin^3D^ cells (top) or in any other conditions (bottom). (H, I) Average ChIP-Seq site profile of cohesin islands detected only in WAPL-degraded cohesin^3D^ cells (H) or in any other conditions. (J) ‘Meta-gene’ analysis showing the distribution of ChIP-Seq signal of SMC1^WT^ and SMC1^3D^ in cells depleted of WAPL or not over an exemplary gene, as indicated (K) Hi-C matrices and ChIP-Seq profiles of chromosome 1 (174-177 Mb) obtained from WAPL degraded cohesin^WT^ and cohesin^3D^ cells. Dashed circles indicate chromosomal contacts at cohesin islands that are enriched following WAPL degradation specifically in cohesin^3D^ cells. (L) Aggregation peak analysis of chromosomal contacts at cohesin islands as in (K). The enhanced and reduced contacts in WAPL depleted cohesin^3D^ cells relative to those in WAPL depleted cohesin^WT^ cells are shown in red and blue respectively.

By contrast, SMC1^3D^ was only enriched in 18,997 peaks (Figure 5A-E), which on average only contained 50% of the read counts found in SMC1^WT^ peaks (Figure S4C). Most of these SMC1^3D^ peaks overlapped in their genomic positions with SMC1^WT^ peaks (95%; Figure S4B). WAPL degradation increased the amount of cohesin^3D^ on chromatin (Figure 5F, compare SMC1, SMC3 and SCC1 signals in lanes 7 and 8) and accordingly the number of SMC1^3D^ ChIP-Seq peaks (to 32,329; Figures 5B and S4B) and their read counts (Figure S4C), but not to the same extent as for SMC1^WT^ (Figures 5B and S4C).

More detailed analyses of these data revealed two interesting features. First, SMC1^3D^ peaks responded differently to WAPL depletion depending on whether they overlapped with CTCF sites or not. Whereas WAPL degradation increased SMC1^3D^ ChIP-Seq signals at CTCF sites only to levels observed for SMC1^WT^ peaks in the presence of WAPL (Figure 5C), SMC1^3D^ signals at non-CTCF sites were increased to levels observed for SMC1^WT^ in the absence of WAPL (Figure 5D). In other words, WAPL depletion allowed SMC1^3D^ ChIP-Seq signals to ‘catch up’ to those of SMC1^WT^ at non-CTCF sites but not at CTCF sites. We suspect that this differential response of SMC1^3D^ ChIP-Seq signals to WAPL depletion is caused by a partial deficiency of cohesin^3D^ in accumulating at CTCF sites. Consistent with this possibility we found that fewer SMC1^3D^ ChIP-Seq peaks overlapped with CTCF sites in both the presence and absence of WAPL than SMC1^WT^ peaks (Figures 5E). Second, in WAPL depleted cells only 78% of SMC1^3D^ peaks overlapped with SMC1^WT^ peaks (as opposed to 95% in the presence of WAPL; Figure S5B) and some of these SMC1^3D^ specific ChIP-Seq signals were broader than canonical cohesin peaks (Figure 5A). Among all the conditions, we identified 1,095 such broad ‘cohesin islands’, which occupied more than 5 kb, in contrast to canonical cohesin ChIP-Seq peaks, which cover regions of ∼0.3-0.6 kb. Of these cohesin islands, 756 were only detectable in cohesin^3D^ cells after WAPL degradation (Figure 5G and H) and almost half of these were found near the 3’ ends of genes (Figure 5G), whereas only 25 % of the remaining 339 were found there (Figure 5G and I). To test whether SMC1^3D^ became generally more enriched in the 3’ regions of genes than SMC1^WT^ we plotted their ChIP-Seq signals onto all genes after aligning these independently of their absolute length at their transcription start sites (TSSs) and transcription termination sites (TTSs). This ‘meta-gene’ analysis revealed that SMC1^WT^ signals were higher than SMC1^3D^ signals both before and after WAPL degradation at TSSs, whereas the opposite was seen at TTSs where WAPL depletion increased SMC1^3D^ signals to levels much above those of SMC1^WT^ (Figure 5J). Consistent with this, 509 out of 757 cohesin islands which were specifically detected in WAPL degraded cohesin^3D^ cells were found at convergently transcribed gene ends. The accumulation of SMC1^3D^ at the 3’-ends of genes coincided with the enhancement of existing long-range *cis*-interactions or the appearance of new *cis*-interactions that were anchored at these sites (Figure 5K and L). The accumulation of SMC1^3D^ at these sites is reminiscent of the accumulation of wild-type cohesin in ‘cohesin islands’ at the 3’-ends of convergently transcribed genes in cells depleted of WAPL and CTCF (Busslinger et al., 2017), except that SMC1^3D^ displayed such a behavior after depletion of WAPL alone, i.e. in the presence of CTCF. With respect to the formation of cohesin islands, the behavior of cohesin^3D^ therefore resembles that of CTCF depletion.

## Discussion

### Cohesin’s cohesion and loop extrusion functions can be separated

It has long been suspected that cohesin differs from other members of the SMC complex family in that it is able to connect DNA molecules in both *trans* (Guacci et al., 1997; Michaelis et al., 1997) and *cis* (Hadjur et al., 2009; Nativio et al., 2009; Wendt and Peters, 2009; Wendt et al., 2008), whereas other SMC complexes are only known to form *cis*-loops (Davidson and Peters, 2021). It has been proposed that cohesin forms *trans*-contacts by entrapping replicated DNA molecules (Haering et al., 2002) and *cis*-contacts by extruding single DNA molecules (Nasmyth, 2001; Sanborn et al., 2015, Fudenberg et al., 2016; Nichols and Corces, 2018). There is now direct experimental evidence for these hypotheses (Davidson et al., 2019; Haering et al., 2008; Kim et al., 2019) but the mechanistic basis of these two functions remains poorly understood. It is therefore unknown whether cohesin uses similar or distinct mechanisms to generate cohesion and to perform loop extrusion.

Given that most proteins, which have multiple cellular functions, perform these via one and the same mechanism, it would be plausible to assume that cohesin’s cohesion and loop extrusion functions are also based on the same or closely related mechanisms. However, our finding that these functions can be separated indicates that this is not the case. Instead, our results, combined with the proposals that cohesin entraps replicated DNA molecules to mediate sister chromatid cohesion (Haering et al., 2008) but can perform loop extrusion without having to entrap DNA (Davidson et al., 2019, Pradhan et al., 2021), indicate that cohesin uses distinct mechanisms to perform these two functions. This notion is also supported by the observation that the maintenance of cohesion requires PDS5 proteins and sororin but not NIPBL (Ciosk et al., 2000; Hartman et al., 2000; Nishiyama et al., 2010; Panizza et al., 2000; Rankin et al., 2005; Srinivasan et al., 2019), whereas loop extrusion depends on the continuous presence of NIPBL but does not require PDS5 and sororin (Davidson et al., 2019).

The cohesin mutants, which we characterized here are defective in cohesion but able to extrude DNA, indicating that cohesin’s ability to perform the latter function does not depend on the former. A recent study reported that a mutant of the meiotic cohesin kleisin subunit Rec8 (rec8-F204S) can mediate cohesion but is reduced in its ability to generate or maintain chromatin loops in fission yeast cells (Sakuno et al., 2022). Although it has not been demonstrated yet that Rec8 containing cohesin complexes can extrude DNA and that this function is compromised in the rec8-F204S mutant, these observations imply that cohesin’s ability to mediate cohesion does also not depend on its loop extrusion function.

### How do mutations in cohesin’s hinge cause defects in cohesion?

Our observation that WAPL depletion stabilizes cohesin^3D^ on chromatin but does still not enable it to mediate cohesion indicates that the cohesin^3D^ mutant is not simply defective in maintaining cohesion but might be unable to establish cohesion in the first place. According to this hypothesis, the reduced SMC3 acetylation, sororin binding and interactions with PDS5 proteins of cohesin^3D^ and its short chromatin residence time in G2 might be consequences and not the causes of this cohesion defect. Experimental stabilization of Smc1-Smc3 hinge interactions reduces entry of DNA into the cohesin ring and causes lethality in yeast, suggesting that transient hinge opening generates an entry gate for DNA into the ring (Gruber et al., 2006, Collier and Nasmyth, 2022). Consistent with this possibility, transient hinge opening has been observed by high-speed atomic force microscopy (Bauer et al., 2021). The hinge mutant characterized here could therefore be defective in entrapping DNA. Alternatively, it is possible that cohesin^3D^ can entrap DNA but cannot keep it inside its ring structure because the hinge domain re-opens too frequently or precociously.

### Does cohesin entrap the DNA ‘anchors’ of extruded *cis*-loops at CTCF sites?

Unexpectedly, our results indicate that mutation of the inner pore of cohesin’s hinge does not only affect cohesion establishment but also cohesin’s response to CTCF. These effects are less strong than the effects of cohesin^3D^ on cohesion, but several independent observations indicate that the SMC1^3D^ mutation substantially reduces the ability of extruding cohesin complexes to interact with CTCF. First, fewer SMC1^3D^ than SMC1^3D^ ChIP-Seq peaks overlap with CTCF (Figure 5E) and cohesin^3D^ responds to WAPL depletion differently at CTCF and non-CTCF sites (Figure 5C and D), suggesting that cohesin^3D^ is not retained at CTCF sites as well as cohesin^WT^. Second, in cohesin^3D^ cells fewer Hi-C peaks can be detected than in cohesin^WT^ cells (Figure 4F), resembling one of the phenotypes of CTCF depleted cells, which also container fewer Hi-C peaks (Nora et al., 2017; Wutz *et al*., 2017). Third, in WAPL depleted cells cohesin^3D^ accumulates in cohesin islands at the 3’ end of genes (Figure 5A, G and J), whereas wild-type cohesin displays such a behavior only after simultaneous depletion of WAPL and CTCF (Busslinger et al., 2017). In the first two cases, the SMC1^3D^ mutation only partially phenocopies the effects of CTCF depletion, which causes even stronger reductions in cohesin-CTCF co-localization and Hi-C peaks (Busslinger et al., 2017; Nora et al., 2017; Wutz et al., 2017). However, the SMC1^3D^ mutation has a comparably strong effect on the formation of cohesin islands in WAPL depleted cells as WAPL-CTCF co-depletion has on wild-type cohesin (Busslinger et al., 2017). Furthermore, the accumulation of cohesin^3D^ in cohesin islands represents a gain-of-function phenotype, which cannot be explained by the reduced chromatin levels of this mutant and must therefore be caused by a defective response of cohesin^3D^ to CTCF boundaries.

What could this defect be and how could the same SMC1^3D^ mutation affect both cohesion and cohesin-CTCF interactions? One possibility would be that the same amino acid residues in the inner pore of the hinge are independently required for both these processes. However, a simpler and thus more plausible interpretation is that the SMC1^3D^ mutation affects a property of cohesin that is required for both cohesion establishment and cohesin’s response to CTCF. If the inner-pore hinge mutants characterized here abrogate cohesion because they prevent entrapment of replicated DNA molecules inside cohesin’s ring, DNA entrapment via the same route might also be required for cohesin’s proper response to CTCF. According to this hypothesis, cohesin could extrude DNA without entrapping it inside its ring structure (Davidson et al., 2019) but upon encountering CTCF would transiently open the hinge to entrap the ‘anchor’ DNA segments of an extruded loop. Such a mechanism has recently been independently proposed based on the observation that relatively more cohesin complexes can be released by proteolytic cleavage of their kleisin subunits from CTCF sites than from other chromatin regions (Li and Dekker, 2021). Although more work will be required to test this hypothesis, it is attractive as it could not only explain some of our observations and those of Li and Dekker (2021) but also how extruded loops can be maintained for prolonged periods of time (Vian et al., 2018) and why some loop anchoring cohesin complexes have residence times on chromatin that are as long as those of cohesive cohesin complexes, which are thought to entrap DNA (Wutz et al., 2020).

## Acknowledgements

We thank the staff at the IMBA/IMP/GMI BioOptics, Molecular Biology Service and Next-generation sequencing for technical support; Georg Winter for providing dTAG7; Iain Patten for useful comments on the first draft; Madhusuhan Srinivasan and Kim Nasmyth for helpful discussions and critical reading of the manuscript and all the Peters lab members for discussions. K.N was supported by an EMBO long-term fellowship (ALTF 1335 - 2016) and a HFSP Long-term fellowship (LT001527/2017). Research in the laboratory of D.W.G. is supported by the Austrian Academy of Sciences, the Vienna Science and Technology Fund (WWTF; projects LS17-003 and LS19-001), and the European Research Council (ERC) under the European Union’s Horizon 2020 research and innovation programme (grant agreement no. 101019039). Research in the laboratory of J.-M.P is supported by Boehringer Ingelheim, the Austrian Research Promotion Agency (Headquarter grant FFG-852936), the European Research Council under the European Union’s Horizon 2020 Research and Innovation Programme (1020558), the Human Frontier Science Program (RGP0057/2018), and the Vienna Science and Technology Fund (LS19-029). J.-M.P. is also an adjunct professor at the Medical University of Vienna.

## Author contribution

K.N. designed research, generated constructs, and cell lines, performed and analyzed cellular experiments, and wrote the manuscript. I.F.D performed and analyzed loop extrusion experiments. R.S. analyzed ChIP-Seq and Hi-C data. W.T. performed calibrated ChIP-Seq experiment. G.W. generated Hi-C libraries. P.B. performed, analyzed scsHi-C experiments and wrote a script for image analysis. G.L. and M.P. performed and analyzed ATPase assays. A.S. performed protein sequence analysis. D.W.G. supervised scs-HiC experiments. J.-M.P. supervised research and wrote and finalized the manuscript with input from all authors.

## Declaration of interests

The authors declare no competing interests.

## Material and Methods

### Antibodies

The following in-house generated rabbit polyclonal antibodies were used: SMC1 (ID:A1027), Sororin (ID:A953), WAPL (ID:A1017) and PDS5B (ID:A770). The following commercially available antibodies were used: SMC3 (Bethyl Laboratories, A300-060A), SCC1 (EMD Milipore corporation, 53A303), PDS5A (Bethyl Laboratories, A300-089A), a-tubulin (Sigma-Aldrich, T5168), Myc (Sigma-Aldrich, 05-724) and histone H2B (Abcam, ab52599). Mouse monoclonal anti-acetyl-SMC3 antibodies were a generous gift from Katsuhiko Shirahige (Nishiyama, 2010). ChIP-Seq was performed Anti-Flag antibodies (Cell signalling, D6W5B).

### Cell culture, generation of CRISPR engineered cell lines, lentivirus transduction and RNAi

HeLa Kyoto cells were cultured in DMEM supplemented with 10% FCS, 0.2 mM L-glutamine and antibiotics. SCC1-Halo-P2A-Tir1, SMC1-AID-mKate2 and Blasticidin-P2A-FKBP^F36V^-WAPL cell lines tagged at the respective endogenous loci were generated by CRISPR/Cas9-mediated homologous recombination using a double-nickase strategy (Ran et al., 2013). Homology arms surrounding the start or stop codons of SCC1, SMC1 and WAPL were cloned into the vector pBSKII. Halo-P2A-Tir1, AID-mKate2 and Blasticidin-P2A-FKBP12^F36V^ coding sequences were introduced before the stop or after the start codon. These homology-directed recombination donor plasmids were transfected to parental cells with pairs of CRISPR guide RNAs which are targeted at N or C terminal of target genes. Seven days after transfection, Halo^TMR^ and mKate2 positive single cells or blasticidin (5 μg/ml) resistant single cells were isolated by FACS sorting or after blasticidin treatment, respectively. Homozygous targeting of genomic alleles was confirmed by genomic PCR and western blotting. To generate cell lines that homogeneously expressed SMC1^WT^ or SMC1^3D^ proteins, lentiviruses were generated by transfecting pRRL-TRE3G vectors encoding SMC1^WT^-Flag or SMC1^3D^-Flag together with the lentiviral packaging vectors pCMVR8.74 (addgene #22036) and pMD2.G (addgene #12259) to LentiX cells. After 72 hours, lentiviral culture medium was harvested and added to HeLa Kyoto cell lines. After infection for 24 hours, puromycin (2 μg/ml) resistant single cells were isolated. For RNAi depletion of Sororin, cells were treated with 30 nM sororin targeting siRNAs with RNAiMax (Invitrogen) for 24 hours, as described previously (Schmitz et al., 2007).

### Live cell imaging, pciFRAP and curve fitting

Cells were seeded on chambered coverglass (Nunc 155409) for 2 days in DMEM supplemented with 10% FCS, 0.2 mM L-glutamine antibiotics and 1 μg/ml doxycycline to induce expression of SMC1^WT^-Flag or SMC1^3D^-Flag. Before imaging SCC1-Halo, cells were incubated with 250 μM HaloTag TMR ligand (Promega, G8251) for 20 min. After washing with pre-warmed cell culture medium three times, cells were incubated for 30 min and then exchanged for pre-warmed phenol red free medium supplemented with 10% FCS, 0.2 mM L-glutamine antibiotics for imaging. For DNA labeling, cells were incubated with 0.5 μM SiR-DNA (Spirochrome), for 2 hours or 0.1 μg/ml Hoechst33342 (SIGMA-ALDRICH) for 20 min before imaging. Live cell imaging was performed using an LSM880 confocal microscope (Carl Zeiss), equipped with a 40x /1.4 numerical aperture (N/A) oil DIC Plan-Apochromat objective at 37 °C and 5 % CO_2_. For quantification of vermicelli, cells were imaged after the addition of dTAG7 (1 μM, a generous gift from Georg Winer) for 5 hours. For pciFRAP, HaloTag ligand building blocks (Succinimidyl Ester (O2) Ligand, Promega) were mixed with 1 M Tris-HCl (pH 8.0), incubated for 60 min at room temperature and added to imaging medium (final 100 μM), following the previously reported protocol (Yamaguchi et al., 2009). Photobleaching half of the nucleus was performed by 2 iterations of a 561 nm diode laser at max intensity after the acquisition of two images. After the bleaching, images were acquired at 1 min intervals for 2 hours. The fluorescent recovery kinetics were quantified using the ZEN2011 software, as the difference between the mean fluorescence signal intensity of the bleached and the unbleached regions followed by background subtraction. pciFRAP curves were normalised to the mean of the pre-bleach fluorescent intensity and to the first image after photobleaching. G1 and G2 phase cells were identified by nuclear or cytoplasmic localization of DHB-mVenus signals, respectively (Spencer et al., 2013). iFRAP curves were fitted using the single exponential function f(t)=EXP(-kOff1*t) or the double exponential function f(t)=a*EXP(-kOff1*t)+(1-a)*EXP(-kOff2*t) in R using the mini-pack.lm package (version 1.2.1) as previously described (Holzmann et al., 2019). Double exponential curve fitting was performed under the constraints that 1/kOFF1 (the dynamic residence time) and 1/kOFF2 (the stable residence time) were in the range of 1-40 min and 5-15 hours, respectively.

### Vermicelli quantification using coefficient of variation measurements

Coefficient of variation measurements were calculated using a custom Python script. In brief, the centremost image within each Z-stack was calculated by first identifying the image with the highest mean pixel intensity in the DNA (Sir-Hoechst) channel. This image was then thresholded to create a binary mask, and the mask was applied to the images at this Z position in the Scc1 and DNA channels. The mean and standard deviation of the pixel values within this mask was then calculated in both channels. The coefficient of variation for the Scc1 channel was subsequently calculated by dividing the standard deviation by the mean.

### Chromatin fractionation and immunoprecipitation

For chromatin fractionation, cell pellets were extracted in a buffer consisting of 20 mM Tris (pH 7.5), 150 mM NaCl, 5 mM MgCl_2_, 2 mM NaF, 10% glycerol, 0.2% NP40, 20 mM b-glycerophosphate, 0.5 mM DTT, and protease inhibitor cocktail (Complete EDTA-free, Roche). Chromatin pellets were separated from whole cell extracts by centrifugation at 2,000 g for 5 min and were washed three times with the same buffer. Genomic DNAs in the resulting chromatin pellets and in whole cell extracts were digested in the same buffer supplemented with Benzonase nuclease (homemade). For immunoprecipitation, cells were lysed in a buffer consisting of 20 mM Tris (pH 7.5), 150 mM NaCl, 5 mM MgCl_2_, 2 mM NaF, 10% glycerol, 0.2% NP40, 20 mM b-glycerophosphate, 0.5 mM DTT, protease inhibitor cocktail (Complete EDTA-free, Roche) and Benzonase nuclease, and incubated on ice for 20 min. After centrifugation at 15,000 rpm for 30 min at 4 °C, supernatants were incubated with anti-Flag M2 affinity beads (SIGMA-ALDRICH) for 2 hours at 4 °C. Beads were washed three times with the same buffer and resuspended in SDS sample buffer.

### Cell synchronisation, chromosome spreading and cell proliferation

To analyse the consequences of WAPL degradation in cohesin^3D^ cells, cells were synchronised at early S phase by two consecutive rounds of treatment with 2 mM thymidine (Sigma) in the presence of 1 μg/ml doxycycline. Two hours prior to release, cells were treated with 200 μM auxin (Andole-3-Acetic Acid, Gold Biotechnology) and dTAG7 were added. After washing with pre-warmed medium, cells were incubated with medium containing 200 μM auxin and 1 μM dTAG7 for 6 hours and harvested for immunoprecipitation. For chromosome spreading, 7 hours after release, cells were treated with 150 ng/ml nocodazole and incubated for 1 hour. Mitotic cells were harvested by mitotic shake off and processed for chromosome spreading and Giemsa staining as previously described with minor modifications (Hauf et al., 2003). Cells were hypotonically swollen in 40 % PBS/60 % tap water for 5 min at room temperature and fixed with freshly made carnoy’s solution (75 % methanol, 25 % acetic acid). Cells were washed with carnoy’s solution several times and were dropped onto glass microscopy slides and dried at room temperature. Slides were stained with 5 % giemsa solution at pH 6.8 for 5 min, washed with tap water, dried and imaged. For scsHi-C, cells were synchronised in early S phase by two consecutive rounds of treatment with 2 mM thymidine and 3 ug/ml (Sigma) aphidicolin, then treated with 2 mM 4sT (Carbosynth) before the release. Cell proliferation was assessed by staining cells with crystal violet (Sigma-Aldrich) 5 days after addition of 200 uM auxin.

### Calibrated ChIP-Seq

An equal number of cells from each condition were mixed with 3 % MEFs expressing a Flag-tagged chromosome-associated protein and then fixed with 1 % formaldehyde for 10 min at room temperature. After quenching by 125 mM Tris-HCl pH 7.5, cells were washed with PBS and snap frozen in liquid nitrogen. Subsequent ChIP-seq experiment procedures were performed as described (Ladurner *et al*., 2016). Briefly, cell pellets were thawed on ice and lysed with lysis buffer (50 mM Tris-HCi pH 8.0, 10 mM EDTA pH 8.0, 1 % SDS, 1 mM PMSF and protease inhibitor cocktail). Cell lysates were sonicated for 4 cycles (30 sec on/off) at maximum power using a Biorupter to shear the genomic DNA. 10x volumes of dilution buffer (50 mM Tris-HCi pH 8.0, 10 mM EDTA pH 8.0, 1 % Triton X-100, 1 mM PMSF and proteasome inhibitorcock tail) were added to lysate. After pre-clearing by incubation with Affi-Prep protein A beads (BioRad, #1560005), the lysates were incubated with Flag antibodies overnight at 4 °C and incubated with the same beads for 3 hours at 4 °C. The beads were washed 2 times with wash buffer 1 (20 mM Tris-HCi pH 8.0, 2 mM EDTA pH 8.0, 1 % Triton X-100, 150 mM NaCl, 0.1 % SDS and 1 mM PMSF), wash buffer 2 (20 mM Tris-HCi pH 8.0, 2 mM EDTA pH 8.0, 1 % Triton X-100, 500 mM NaCl, 0.1 % SDS and 1 mM PMSF), wash buffer 3 (10 mM Tris-HCi pH 8.0, 2 mM EDTA pH 8.0, 250 mM LiCl, 0.5 % NP-40, 0.5 % deoxycholate) and TE buffer (10 mM Tris-HCl pH 8.0, 1 mM EDTA pH 8.0), and then eluted with elution buffer (25 mM Tris-HCi pH 7.5, 5 mM EDTA pH 8.0 and 0.5 % SDS) for 20 min at 65 °C two times. The elutes were treated with RNase-A at 37°C for 1 hour and then proteinase K at 65°C overnight. DNA was purified by phenol/chloroform/isoamyl alcohol (25:24:1) extraction and ethanol precipitation. DNA was resuspended in 100 μl H2O, and ChIP efficiency was quantified by quantitative PCR (qPCR). Sequencing libraries were prepared using the NEBNext® Ultra II kit (NEB, E7645S) following the manufacturer’s instructions, and DNA samples were submitted for Illumina deep sequencing at the Vienna Biocenter Core Facilities.

### ChIP-Seq peak calling and calibration

Prior to treatments and Illumina short read sequencing, a fixed percentage of mouse cells expressing a Flag-tagged marker protein was added to all samples as a calibration reference throughout the entire follow-up processing. Sequencing reads which passed the Illumina quality filtering were mapped against a fusion genome template constructed of mm9 (MGSCv37) and hg19 (GRCh37) reference assemblies. The subsets of uniquely mappable reads from the mm9 resp. hg19 fractions (allowing up to two mismatches each) were separated for further steps. The mm9 fractions from all samples were processed using peak calling by MACS version 1.4.3 with the same fixed shift-size parameters resulting in a genome-wide set of peak regions for the Flag-tagged reference protein in all reference mouse cell populations (Zhang et al., 2008). Considering the expected evenness in signal for all samples in a hypothetical loss-free procedure, out of all peak-sets a union peak set was built with constant coordinates being common in all sample references. This common coordinate set’s read abundances throughout all samples delivers reliable measures for usage in overall calibration of the various treated human hg19 fractions. Peak callings of the actual human (hg19) samples with MACS software versions 1.4 and 2 were applied on full, merged, filtered, and deduplicated replicate data sets using background input (-Dox). As a significance threshold we applied a p-value of 1e-10 resulting in reliable peak sets which are confirmed also through very low numbers of detectable negative peaks in the range of few dozens from -Dox cells. Calibration factors described above were applied to coverage values of alignment profile tracks of the samples to overcome the problem of experimental variation in yield efficiency.

### Calculation of peak overlaps

Genomic overlaps have been calculated using multovl version 1.3 (Aszodi, 2012) and area-proportional threefold Venn diagrams were drawn with eulerAPE (Micallef and Rodgers, 2014). Since occasionally more than one site from one dataset overlaps with a single site in a second dataset, the resulting coordinates of such an overlap contribute to one single entry - a so-called union. Consequently, the overall sum of site counts drops slightly if displayed in union overlaps.

### Identification of cohesin islands

Cohesin islands were identified as areas of clustered peak appearance, as described before (Busslinger et al., 2017). In brief, if individual narrow peaks fulfilling the p-value threshold of 1e-10 are positioned sufficiently close to each other to overlap within the range of half of their own size, then they get merged. If resulting multiply merged peak regions exceed 5Kb in size, then they were categorized as a cohesin island.

### Hi-C data processing

Illumina sequencing was performed on all Hi-C libraries with 150 bp paired-end reads. Two replicate data sets for each library were truncated, filtered, and aligned against the human genome assembly hg19 (GRCh37) using bowtie2 and the HiCUP processing pipeline version 0.7.4, and finally merged (Wingett S, et al. 2015, HiCUP: pipeline for mapping and processing Hi-C data F1000 Research, 4:1310, doi:10.12688/ f1000research.7334.1). Alignments were converted into input for juicer_tools pre (juicer tools version 1.22.01; (Rao et al., 2014) as well as input for HOMER version 4.11 (http://homer.ucsd.edu/homer/; (Heinz et al., 2010). PMID: 20513432). All juicer-based contact matrices used for analysis were Knight-Ruiz normalized. Loop annotation, also known as corner peak annotation, and merging was performed using ‘juicer_tools hiccups’ at default resolution. TAD annotation and insulation scores were generated by the ‘findTADsAndLoops.pl’ script in the HOMER software package. This software scans the relative contact matrices for locally dense regions of contacts or areas with an increased degree of intra-domain interactions relative to surrounding regions. Plots of insulation profiles were made using insulation scores in the bedGraph file format and TAD boundary coordinates from regions around these coordinates. Aggregate peak analysis of hiccups-called Hi-C peaks was performed using ‘juicer_tools apa’. The Hi-C looping peaks detected from all the conditions were classified according to their sizes and used as coordinates for aggregation within Hi-C maps. These aggregated size-classified Hi-C peaks were visualized by plotting the cumulative stack of sub-matrices in a way that Hi-C peaks lie at the center of the matrix and that the resulting APA plot displays the abundance of contacts respective to local contact density.

### scsHi-C library preparation

Sister chromatid-sensitive Hi-C was performed as described previously; full experimental details of the protocol can be found in (Mitter *et al*., 2020; Mitter et al., 2022). In brief, cells were synchronised in G1 phase, before labelling with 4-thio-thymidine (4sT) for one cell cycle. Cells were then synchronised in G2 phase using RO-3306 (Vassilev et al., 2006). Cells were harvested for Hi-C and fixed with 1% formaldehyde. The pellets were then lysed, before digestion of the cross-linked DNA with DpnII, nucleotide fill-in and subsequent ligation. Ligated chromatin was then sheared by sonication and the fragmented chromatin size-selected using AMP Pure XP beads. A biotin pull down was then performed on the size-selected fragments using streptavidin-coated beads before library preparation on the beads using the standard NEBNext Ultra II DNA library prep kit for Illumina (NEB) protocol. The DNA containing solution was then eluted from the beads and precipitated using EtOH precipitation. The precipitated DNA was resuspended in water, before conversion of 4sT to methyl-cytosine using OsO_4_/NH_4_Cl (Mitter *et al*., 2020). The converted DNA libraries were then amplified using qPCR, before a final purification using 0.9x sample volume AMP pure XP beads.

### Cloning, expression and purification of recombinant proteins from insect cells

Human cohesin subunits, SMC1^WT^ or SMC1^3D^, SMC3-Flag, SCC1(TEV)-Halo and 10xHis-STAG1 were assembled into pBig2ab plasmids using biGBax (Weissmann et al., 2016). Flag-Halo-NIPBL-His and MAU2 were tandemlly inserted into pLib as described previously (Davidson et al., 2019). To generate bacmid DNA, pBig2ab and pLib plasmids were transformed into DH10 Multibac bacteria. Bacmids were transfected to Sf9 insect cells by lipofection (FuGENE 6 Transfection Reagent, Promega), to generate baculovirus. After 96 hours incubation at 27 °C, culture medium were harvested as a V0 Baculovirus stock. A 100 ml V1 virus stock was cultured from V0 stock and incubated for 48 hours, and used for protein expression. 1.5 million cells per millilitre SF9 insect cells were infected with V1 Baculovirus, harvested after 48-60 hours incubation at 27 °C, washed with PBS, frozen in liquid nitrogen and stored at -80 °C. Recombinant cohesin complexes and NIPBL-MAU2 were purified via tandem affinity purificaiton using Toyopearl AF-chelate-650M resin (TOSOH Bioscience) and Flag-M2 agarose resin (Sigma, A2220) as previously described (Bauer et al., 2021; Davidson et al., 2019).

### ATPase assay

The rate of ATP hydrolysis by cohesin was measured by using the ADP-Glo Kinase assay (Promega, TM313) as previously described (Panarotto et al., 2022). Recombinant cohesin^WT^ or cohesin^3D^ (25 nM) were incubated with 50 nM NIPBL-MAU2 and 10 ng/μl λ-DNA (NEB, N3011S) in ATP reaction buffer consisting of 20 mM NaH2PO4/Na2HPO4 pH 7.5, 50 mM NaCl, 2.5 mM MgCl2, 1 mM DTT, 0.75 % glycerol, 7.5 mM imidazole, 0.1 mg/ml BSA, 0.3 mM EDTA, 1 mM ATP, for 1 hour at 37 °C. After depleting the remaining ATP via the addition of the Glo reagent, ADP was converted to ATP by the addition of Kinase Detection Reagent. Luminescence of the resulting solution was measured by using a PheraStar FX plate reader (BMG Lab tech) and PheraStar Mars software. The rate of ATP hydrolysis was calculated using a standard curve obtained from a dilution series of ATP and ADP, as described in the ADP-Glo Kinase Assay manual.

### Loop extrusion assay

Perpendicular flow loop extrusion assays were performed essentially as described (Bauer et al., 2021; Davidson et al., 2019). Flow cells were incubated with 1 mg/ml Avidin DN (Vector Laboratories) for 15 min and washed extensively with DNA buffer (20 mM Tris pH 7.5, 150 mM NaCl, 0.25 mg/ml BSA (ThermoFisher Scientific; AM2616)). 40 μl of end-biotinylated λ-DNA (Davidson *et al*., 2016) was diluted to 1 pM in DNA buffer supplemented with 20 nM Sytox Orange (ThermoFisher Scientific; S11368) and introduced into flow cells at 5 *μ*l/ min. Flow cells were washed with 20 *μ*l of wash buffer 1 (50 mM Tris pH 7.5, 200 mM NaCl, 1 mM MgCl2, 5% glycerol, 1 mM DTT, 0.25 mg/ml BSA, 20 nM Sytox Orange) at 5 *μ*l/min. Flow was then switched to perpendicular mode and a further 350 *μ*l of wash buffer 1 was introduced at 100 *μ*l/min. 400 *μ*l of wash buffer 2 (50 mM Tris pH 7.5, 50 mM NaCl, 2.5 mM MgCl_2_, 0.25 mg/ml BSA, 0.05% Tween-20, 20 nM Sytox Orange) was then introduced at 100 *μ*l/min, followed by 100 *μ*l of imaging buffer (50 mM Tris pH 7.5, 50 mM NaCl, 2.5 mM MgCl_2_, 0.25 mg/ml BSA, 0.05% Tween-20, 0.2 mg/ml glucose oxidase (Sigma; G2133), 35 mg/ml catalase (Sigma; C-40), 9 mg/ml b-D-glucose, 2 mM Trolox (Cayman Chemical; 10011659)) and 5 mM ATP (Jena Biosciences; NU-1010-SOL)) supplemented with 220 nM Sytox Orange at 100 *μ*l/min. Stock solutions of glucose oxidase, catalase and glucose were prepared as described (Bauer *et al*., 2021). Recombinant wild type or 3D cohesin was then introduced into flow cells at 1.5 nM final concentration together with recombinant NIPBL-MAU2 at 7 nM final concentration in 250 *μ*l imaging buffer supplemented with 220 nM Sytox Orange at 30 *μ*l/min. Loop extrusion experiments were performed at 37 °C. Time-lapse microscopy images were acquired using a Zeiss TIRF 3 Axio Observer setup equipped a with 561 nm laser, an Andor iXon Ultra 888 camera and an Alpha Plan-Apochromat 100x/1.46 DIC oil objective. 100 ms exposure time images were acquired at 4 s intervals. Image analysis was performed in ImageJ. To determine the fraction of DNA molecules that were extruded into loops, the number of doubly tethered DNAs that formed loops during the 8 min 20 s cohesin flow-in time period was divided by the total number of doubly tethered DNAs that were stretched by perpendicular flow. DNAs that were singly-tethered or oriented in such a way that loops were obscured were excluded from analysis. To determine the rate of LE, a custom ImageJ script was used to measure the length of DNA molecules before LE and the length of DNA not contained within the loop during LE. These measurements were converted into kbp, plotted as a function of time in Prism 9 (Graphpad) and used to calculate the LE rate.

## Data availability

The Hi-C and ChIP-Seq sequencing data from this publication will be deposited in the Gene Expression Omnibus (GEO).

---

**Figure S1.**
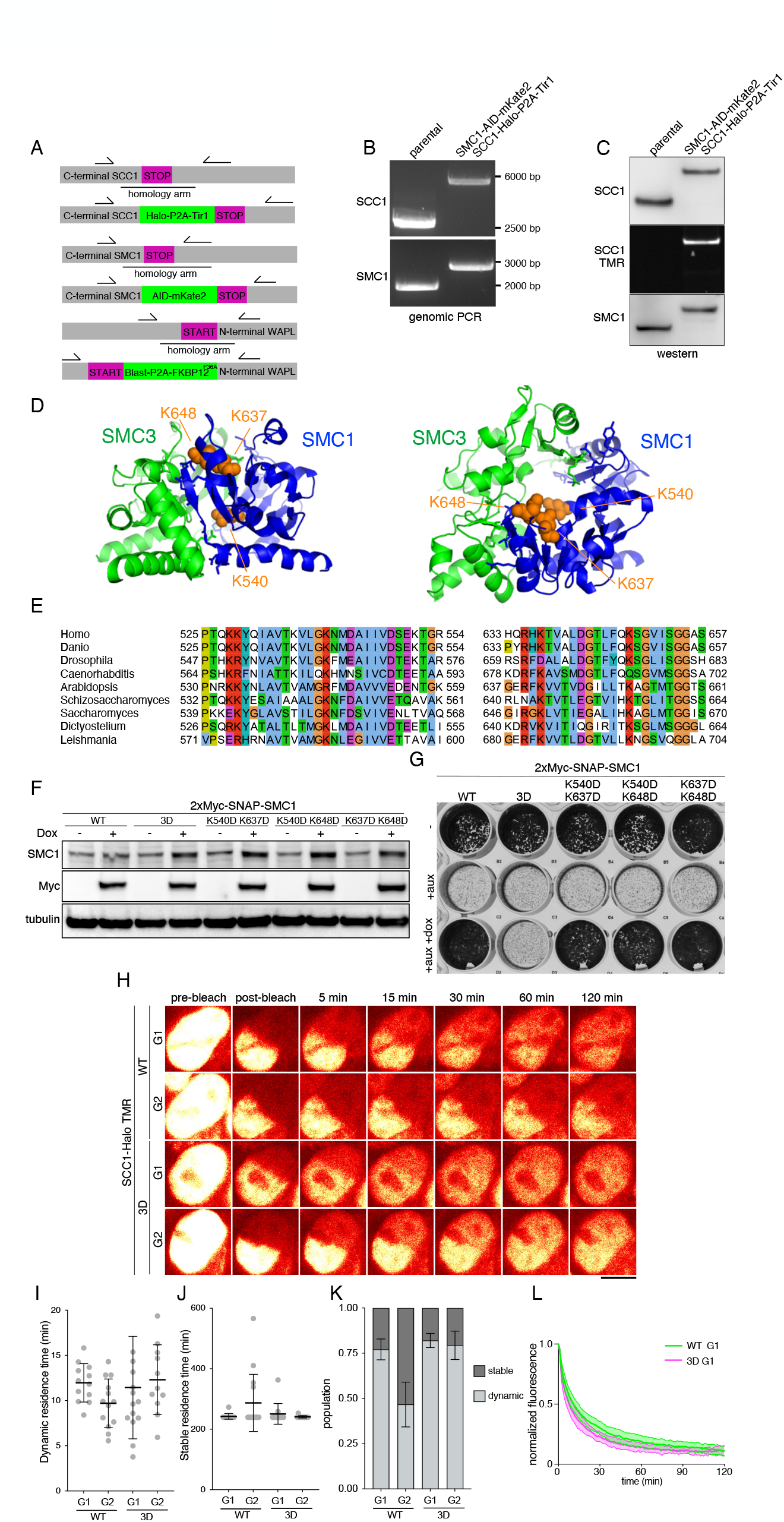
(A) Schematic images of CRISPR engineered AID and FKBP^F36V^ tagging strategy. The Oryza sativa F-box transport inhibitor response-1 auxin receptor protein, Tir1, was expressed following SCC1-Halo translation via T2A self-cleaving peptides. (B, C) Genotype analysis of parental HeLa cells and homozygous SMC1-AID-mKate2/SCC1-Halo-P2A-Tir1 cells by genomic PCR (B) and immunoblotting (C). (D) Structure of the mouse cohesin’s hinge domain and highlighting three lysine residues inside of the hinge domain. (E) Sequence alignment of SMC1 around K540, K637 and K648 residues (human) among different species. (F) Whole cell extracts of cells ectopically expressing Myc-tagged SMC1^WT^, SMC1^540D/K637D^, SMC1^540D/ K648D^, SMC1^K637D/K648D^ and SMC1^540D/K637D/K648D^ were analyzed by immunoblotting as indicated. (G) Images of plates containing indicated cells, analyzed by Crystal Violet staining for the presence of viable cells after addition of auxin for 5 days and subsequent addition of doxycycline for 48 hours. (H) Representative time-lapse images of SCC1-Halo^TMR^ after photobleaching in cohesin^WT^ and cohesin^3D^ cells in G1 and G2-phase. (I,J) Quantification of dynamic (I) and stable (J) residence time of SCC1-Halo^TMR^ in cohesin^WT^ and cohesin^3D^ cells in G1 and G2-phase. (K) Quantification of stably and dynamically chromosomal bound populations of SCC1-Halo^TMR^ in cohesin^WT^ and cohesin^3D^ cells in G1 and G2-phase. (L) Normalized recovery kinetics of SCC1-Halo signals after photobleaching of cohesin^WT^ and cohesin^3D^ cells in G1-phase.

**Figure S2.**
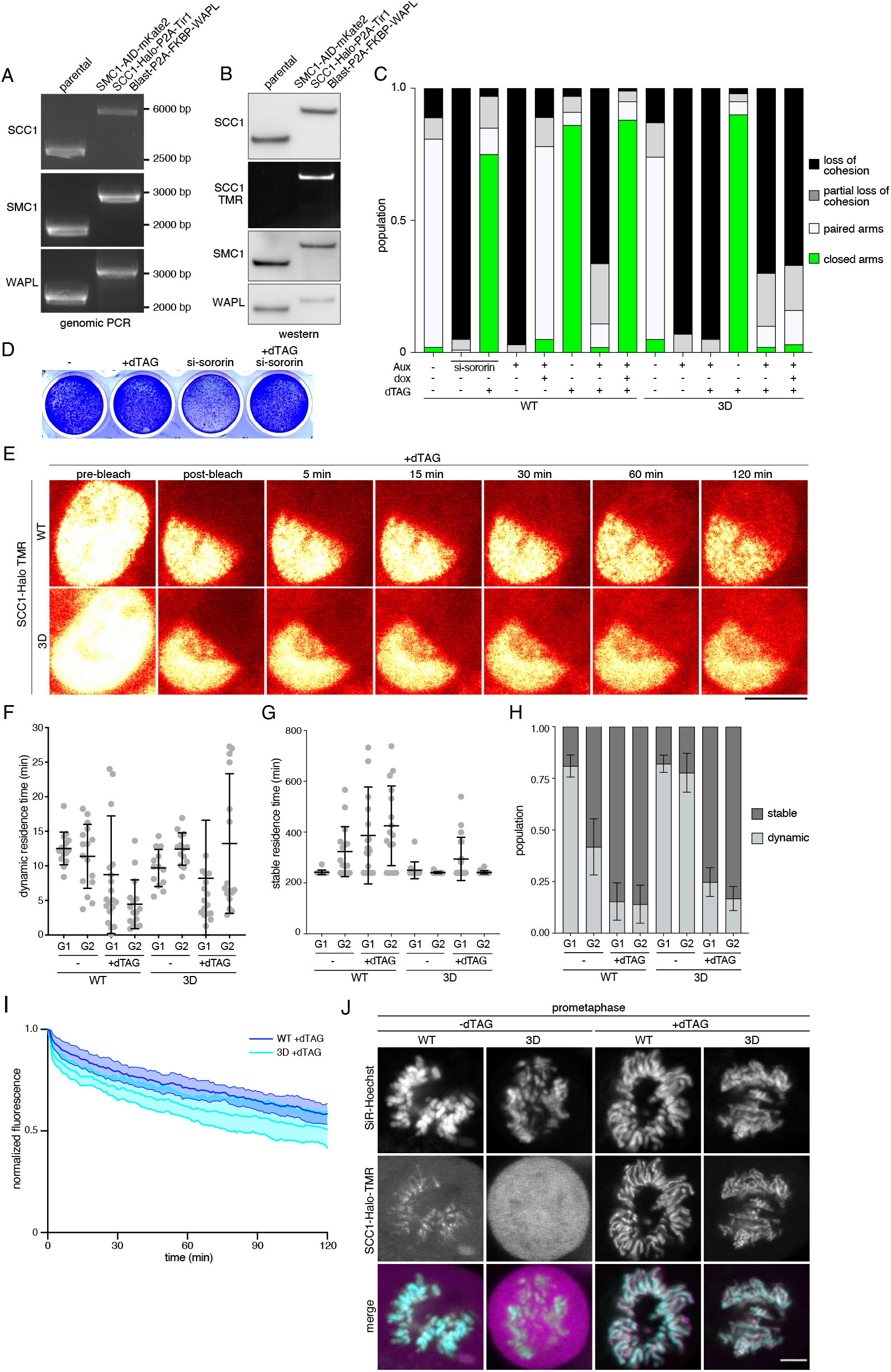
(A, B) Genotype analysis of parental HeLa cells and SMC1-AID-mKate2/SCC1-Halo-P2A-Tir1/ FKBP^F36V^ WAPL cells by genomic PCR (A) and immunoblotting (B). (C) All the quantification of the cohesion phenotypes shown in Figure 2C. (D) Images of plates containing indicated cells, analyzed by Crystal Violet staining for the presence of viable cells after depletion of Sororin by RNAi for 72 hours and subsequent addition of dTAG7 or not for 5 hours. (E) Representative SCC1-Halo^TMR^ images after photobleaching in cohesin^WT^ and cohesin^3D^ cells following WAPL degradation in G1-phase. (F, G) Quantification of dynamic (F) and stable (G) residence time of SCC1-Halo^TMR^ in cohesin^WT^ and cohesin^3D^ cells in G1 and G2 phase following degradation of WAPL or not. (H) Quantification of stably and dynamically bound populations of SCC1-Halo^TMR^. (I) Normalised recovery kinetics of SCC1-Halo^TMR^ after photobleaching of cohesin^WT^ and cohesin^3D^ cells following degradation of WAPL in G1 phase. (J) Live cell imaging of SCC1-Halo^TMR^ in mitotic cohesin^WT^ and cohesin^3D^ cells from which WAPL had been depleted or not. DNA was counterstained with SiR-DNA. Scale bar, 5 μm.

**Figure S3.**
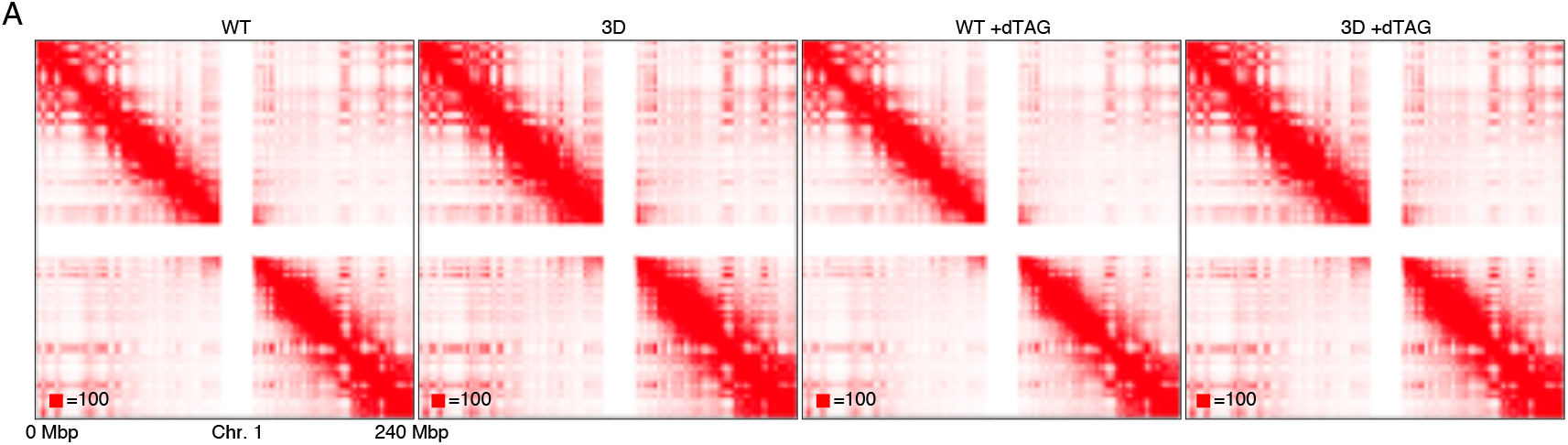
(A) Coverage-corrected Hi-C contact matrices of entire chromosome 1.

**Figure S4.**
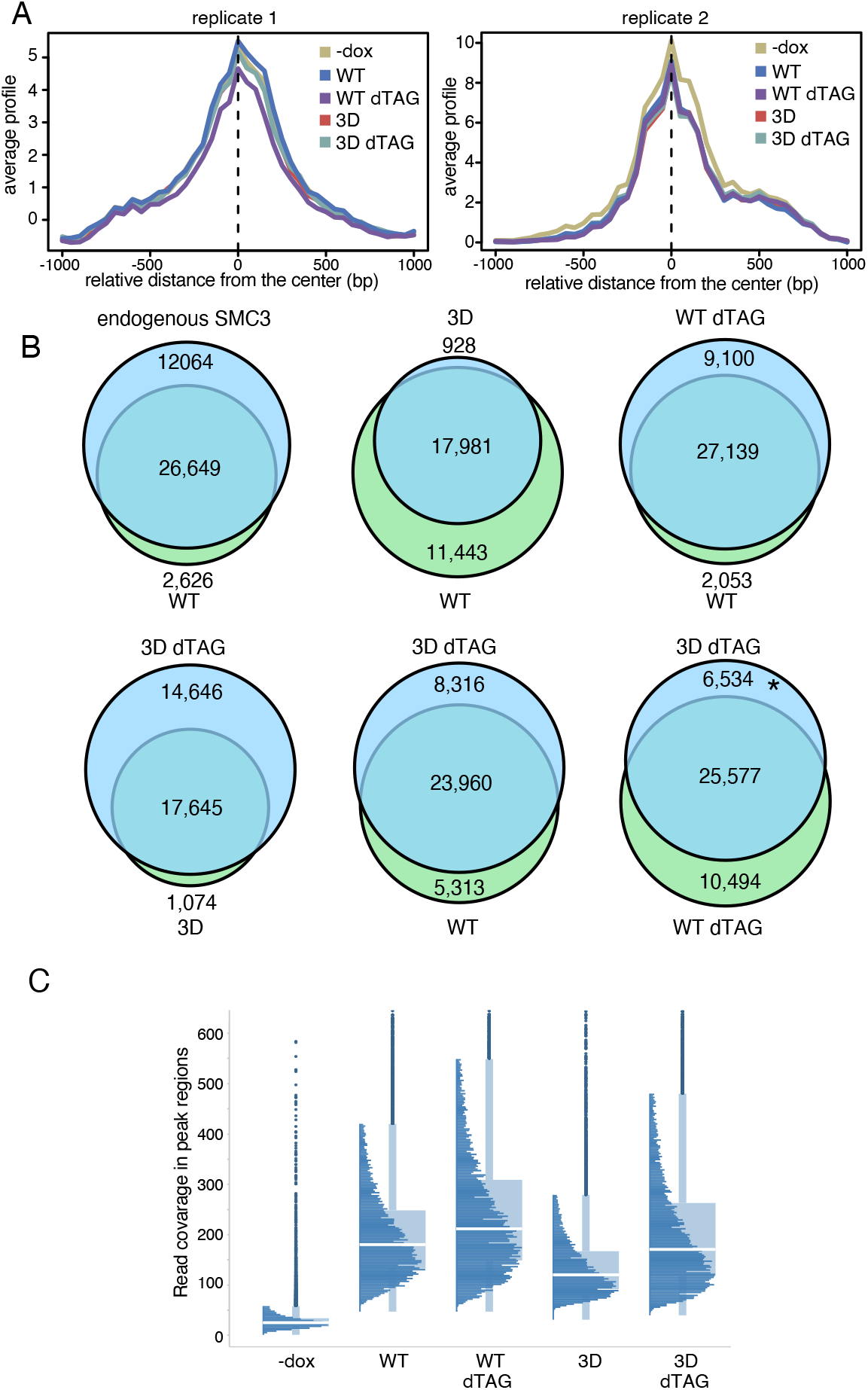
(A) ChIP-Seq signal density of Flag-tagged marker proteins peaks at their detected locations along the mouse genome sequences by peak calling from replicate 1 and replicate 2. (B) Venn diagrams representing genome-wide co-localisation of ChIP-Seq peaks between indicated conditions. (C) Read counts of samples and input inside genomic coordinates commonly detected as peaks.

## Notes

### Competing Interest Statement

The authors have declared no competing interest.

